# A novel view on the mechanism of biological activity of antifreeze proteins

**DOI:** 10.1101/2021.09.22.461391

**Authors:** Bogdan S. Melnik, Ksenia A. Glukhova, Evgeniya A. Sokolova, Irina V. Balalaeva, Alexei V. Finkelstein

## Abstract

The adaptation of organisms to sub-zero temperatures is an intriguing problem in biology and biotechnology. The ice-binding antifreeze proteins are known to be responsible for the adaptation, but the mechanism of their action is still far from being clear. Here we show that: (i) in contrast to common belief, ice-binding proteins do not reduce the water freezing temperature and even raise (1) the ice melting point; (ii) at sub-zero temperatures (to ≈ -30°C), ice can be formed only on ice-binding surfaces, but, for kinetic reasons, not in bulk water; (iii) living cells have some large surfaces, which can bind the antifreeze proteins. These facts allow suggesting that the task of antifreeze proteins is not to bind to the ice crystals already formed in the cell and stop their growth or rearrangement, but to bind to those cell surfaces where the ice nuclei can form, and thus to prevent ice formation completely.

---

Ice-binding proteins (IBPs), also termed antifreeze proteins (AFPs), are synthesized by various organisms to survive at sub-zero temperatures. These proteins were first found in the blood of fish living in the Arctic and Antarctic waters (2–4). Mechanism of action of these proteins is far not clear (5, 6), but it is commonly believed that IBPs either decrease the freezing point of the bodily fluids by several degrees (7) (note that such a decrease of the freezing point by small molecules, like salts or alcohols, would require 300-500 times more material) or bind to ice crystals and prevent their growth, which is fatal to living tissues and cells (8–10).

In this paper, we show that: (I) ice-binding proteins (IBPs) do not reduce the water freezing point, but, on the contrary, raise (1) the ice melting point; (II) at subzero temperatures (from 0°C to ≈-30°C), ice can be formed only on ice-binding surfaces, but, for kinetic reasons, not in bulk water; seemingly, it is these ice-binding surfaces that should be the targets for proteins with the antifreeze activity; (III) living cells have large surfaces that bind IBPs.

These three facts suggest the mechanism of action of antifreeze proteins: their aim is not to bind to the growing ice crystals and stop their growth, but to bind to those cell surfaces where the ice nuclei can form, and thus to prevent ice formation completely.

It is generally accepted (7, 11) that ice-binding proteins decrease the freezing point of water and biological fluids. However, our experiments made us reconsider this standpoint.

The experiments were carried out with a mutant form of the protein cfAFP described in Refs. (12–14); cfAFP is an antifreeze protein of a spruce budworm (*Choristoneura fumiferana)*, a moth whose larvae winter in needles (11) at a temperature of about -30°C. Because this mutant protein is less susceptible to aggregation during isolation and purification, it is most convenient for experiments. As compared to the wild type cfAFP, the mutant was a little shortened and two of its SS bonds were removed by replacement of four cysteines with valines and alanine (for details, see Ref. (15)). This mutant form retains the ability of ice binding (15); we will call it mIBP83.

Our experiments evidenced no notable change in the freezing temperature of a solution supplemented with mIBP83 (Fig. 1, Table 1), while a slight increase of the ice melting point was observed there (Fig. 2).

**Fig. 1.**
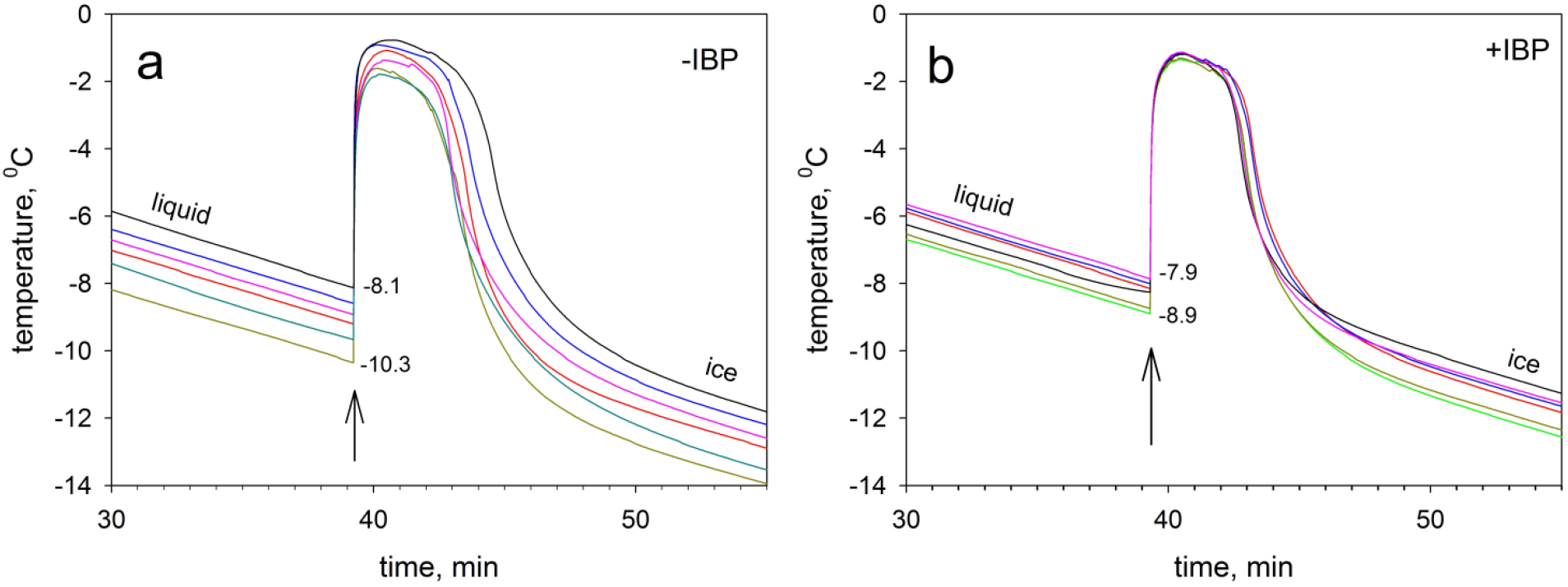
Experiments on freezing solutions. In each individual experiment, a test tube with 150 μl of the solution was cooled from 0°C to -17°C at a rate of -0.24 °C/min. **a**, 20 mM sodium phosphate buffer at pH 7. **b**, a solution of 0.6 mg/ml of mIBP83 in the same buffer. At least 15 independent experiments were performed for each kind of solution. Six experimental curves are shown in each panel as an example. The arrow marks the moment when the solutions freeze. To make it easier to compare the curves, they are aligned at this freezing point. Similar experiments were carried out with distilled water and, as a control, with carbonic anhydrase B (a protein that was never suspected to be antifreeze) in the same phosphate buffer. The ranges of freezing temperatures obtained in the experiments are shown in Table 1.

**Table 1.**
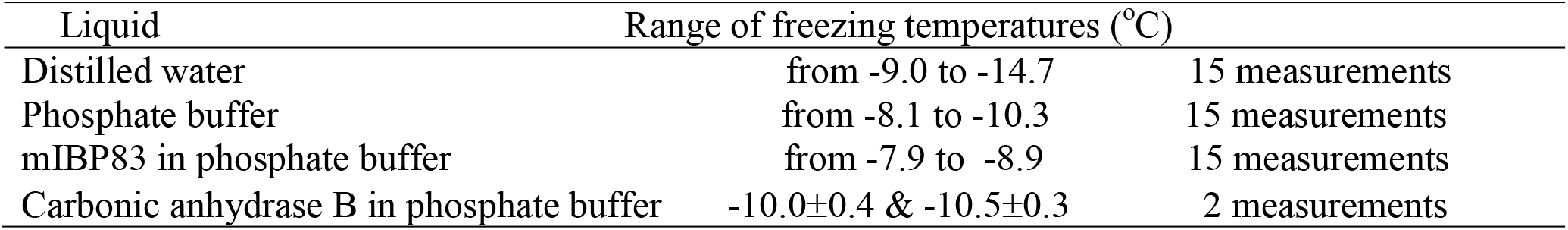
Freezing temperatures for explored liquids

| Liquid | Range of freezing temperatures ($^{\circ}\text{C}$) | |
| --- | --- | --- |
| Distilled water | from $-9.0$ to $-14.7$ | 15 measurements |
| Phosphate buffer | from $-8.1$ to $-10.3$ | 15 measurements |
| mIBP83 in phosphate buffer | from $-7.9$ to $-8.9$ | 15 measurements |
| Carbonic anhydrase B in phosphate buffer | $-10.0 \pm 0.4$ & $-10.5 \pm 0.3$ | 2 measurements |

**Fig. 2.**
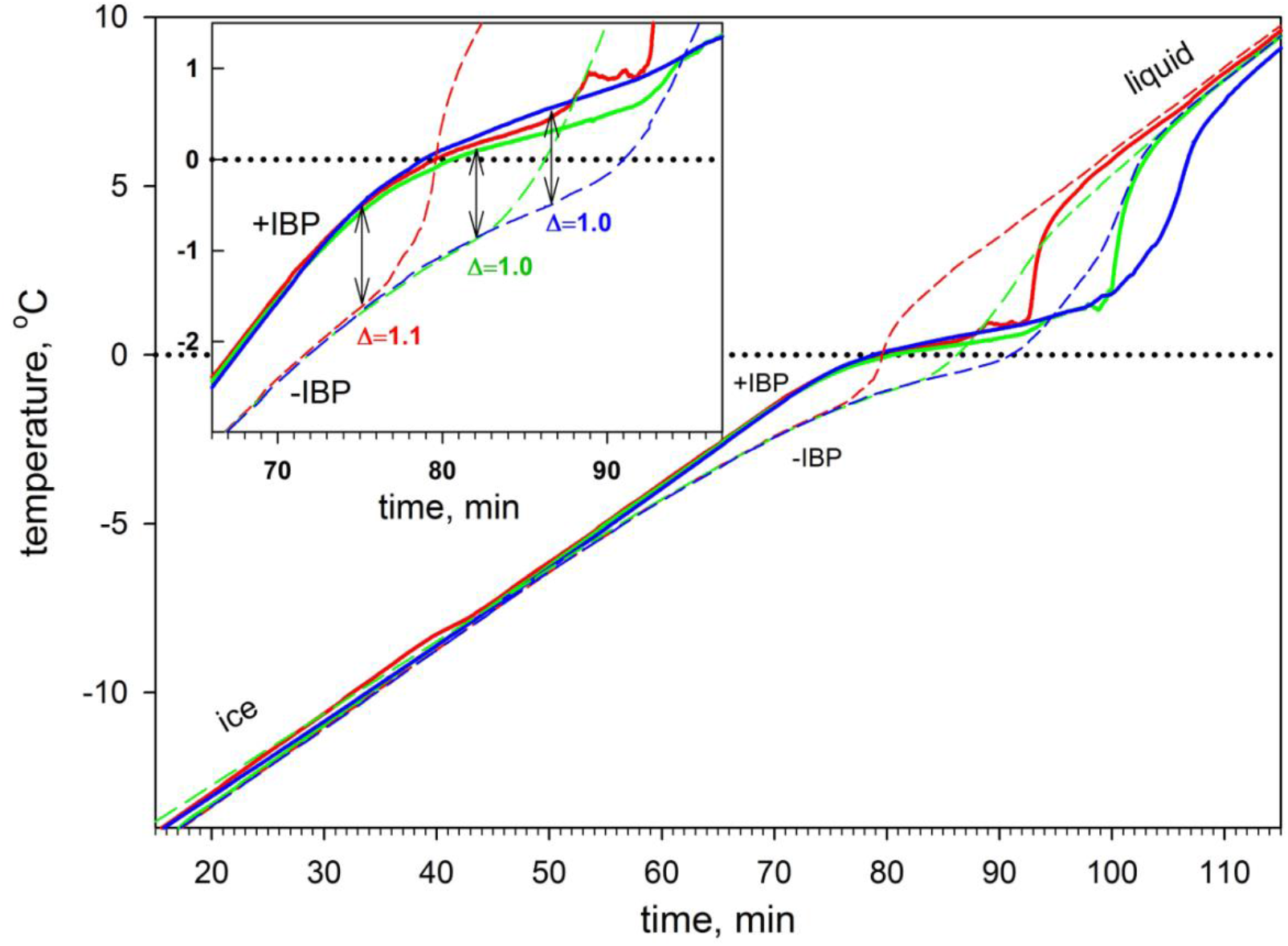
Experiments on melting. In each experiment, a test tube with 100 μl (red lines), 150 μl (green lines) or 200 μl (blue lines) of a frozen solution was heated from -17°C to +10°C at a rate of 0.24 °C/min. Solid lines (+IBP): samples with 0.6 mg/ml of mIBP83 in sodium phosphate buffer (20 mM at pH 7); dashed lines (-IBP): samples containing only the buffer. The central parts of the same curves are shown in the insert at a larger scale. Here, the double arrows show the temperature differences between the +IBP (solid) and -IBP (dashed) lines. The errors in the calculated differences Δ depend mainly on the alignment of the curves (made in the region of -7°C) and do not exceed 0.2°C.

Figure 1 and Table 1 show that freezing of all studied solutions occurs not at 0°C but below -7.9°C (this means, in particular, that blood freezing *per se* cannot be dangerous for the Arctic and Antarctic fishes since the ocean temperature is never below -2°C).

When ice arises in a supercooled liquid explored, the liquid abruptly takes the heat released by the ice; this is detected as a jump in the temperature of the test tube. Then follows a short period with sub-zero temperature when the rest of the liquid freezes, and then we see a rapid decrease in the temperature of the test tube to the temperature of the thermostat.

The phenomenon of supercooling of liquids before freezing is well known (16, 17). Associated with the kinetics of nucleation of crystallization, it will be considered below. We see that the freezing temperature can vary by a few degrees from experiment to experiment performed with different samples of the same liquid. The theory (see below) shows that freezing begins not in bulk liquid but with the appearance of a crystal nucleus at the boundary of the liquid (including at dust particles) so that a smallest scratch at these boundaries can strongly affect the speed and temperature of the appearance of the freezing nucleus. Apparently, this is what makes ice freezing experiments poorly reproducible.

Our ice melting (in contrast to ice freezing) experiments were reproducible with a high accuracy. While the ice is cold and does not melt, its temperature linearly depends on the heating time (Fig. 2). When the melting begins (from the surface – see Ref. (18)), the ice surface absorbs heat, and the temperature in the ice center (where the measuring thermocouple is located) “lags” behind the thermostat temperature rise. The duration of the lag period grows with the increasing volume of the sample (Fig. 2). When the entire ice has melted, the liquid in the test tube quickly heats up and again becomes linearly dependent on the heating time.

Figure 2 shows that ice melting in the presence of an ice-binding antifreeze protein (+IBP) occurs not at the temperature of ice melting in the “pure” (-IBP) buffer but at a higher temperature. The inset in figure 2 shows that the presence of IBP increases the melting point by approximately +1.0°C. It should be noted that superheated (by ≈+1.0°C) ice has been also observed (1) in Antarctic notothenioid fishes: their AFPs inhibit ice melting both *in vivo* and *in vitro*. The phenomenon of a noticeable (for more than +1°C) overheating of crystals before melting is observed, as known, very rarely (16–21), since the surface of the crystal is covered with a thinnest layer of disordered molecules that resembles a supercooled liquid (18, 22) which can serve as the place of rapid nucleation of the liquid phase. The IBP adherence to the surface of a crystal can remove this surface liquid, thereby hindering the nucleation of the liquid phase.

From the experiments described above, as well as from the published data (1), one can conclude that ice-binding proteins do not decrease the temperature of the water-to-ice transition, and, on the contrary, even somewhat increase the temperature of the ice-to-water transition. Therefore, ice-binding proteins in principle cannot work according to the mechanism used by classic low-molecular antifreezes, e.g., alcohols.

To elucidate the mechanism of antifreeze activity of ice-binding proteins, we can turn to the theory of the first order phase transitions (16, 17) describing nucleation of crystals (and specifically of ice) and evaluate the possibility of ice formation in the living cell and bodily fluids under various conditions.

Firstly, we consider the formation of a 3D crystal nucleus in bulk liquid.

The free energy increment upon the formation of a compact piece of a new three-dimensional phase (e.g., a crystal) consisting of *n* particles can be approximately estimated as

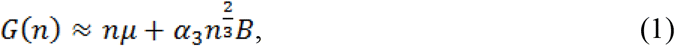

where *μ* < 0 is the chemical potential of a molecule in the “new” phase (e.g., in ice) minus that in the “old” one (e.g., in a liquid), *B* > 0 is the additional free energy of the molecule on the surface of the “new” phase, 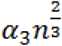 is the number of molecules on the surface of a compact piece of the new phase consisting of *n* particles (for a sphere, 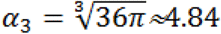; for a cube, *α*_3_ = 6 ; i.e., for our estimates, we can assume that *α*_3_ ≈ 5). The free energy of less compact (with large α) pieces is higher than that of compact ones, which would create a high barrier on a pathway to the new phase; so, considering its nucleation, we may ignore “uncompetitive” slow paths going via non-compact structures.

In a biological object, ice is expected to appear at relatively small negative temperatures. As we will see below, the ice nuclei are rather large at these temperatures; this justifies our neglect of the nucleus structure details.

A little below the point of thermodynamic equilibrium of phases (0°C, i.e., *T*_0_=273°K), ice is more stable than water (i.e., 0 < − *μ*), but only a little, so that − *μ* ≪ *k*_*B*_*T*_0_ (where *k*_*B*_ is the Boltzmann constant), and −*μ* ≪ *B* (while μ≡0 at *T*_0_=273°K). In this case, *G*(*n*) at first grows with increasing *n* (see Eq. (1)), then passes through a maximum of *G*^#^ in the “transition state” (corresponding to the “nucleus” of the ice piece), and then decreases. The maximum of *G*(*n*) is achieved when 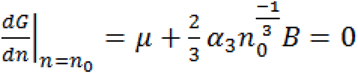, i.e., where the “nucleus” contains

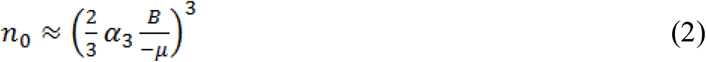

particles; and the “seed”, i.e., the minimally stable piece of ice (with *G*(*n*) = 0 at *n* > 0) contains

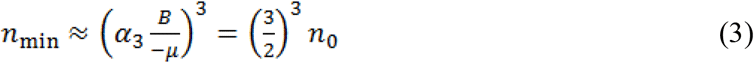

particles. The free energy of the transition state(16, 17)^16,17^ is

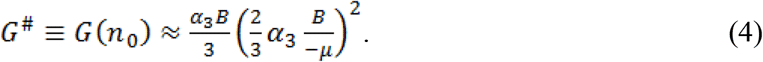

Since *α*_3_≈ 5, then 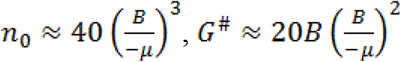.

At the temperature *T*_0_, *μ* ≡ 0 by definition, at *T*_0_ − Δ*T* (where Δ*T* ≪ *T*_0_), 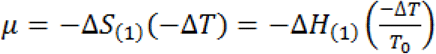 according to the classical equations of thermodynamics (here Δ*S*_(1)_ and Δ*H*_(1)_ are the entropy and enthalpy of water freezing per 1 molecule). The experimental results (see Ref. (23)) show that in water, Δ*H*_(1)_ ≈ −6.0 kJ/mol ≈ −2.6 *k*_*B*_*T*_0_, so that

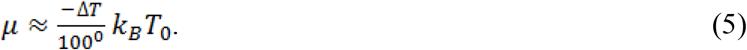

An experimental estimate of *B* (≈ 0.85 *k*_*B*_*T*_0_ near *T*_0_ = 273 К) follows from the free energy of the ice/water interface (24), ≈32 erg/cm^2^, and because an H_2_O molecule occupies ≈ 10Å^2^ of the interface *B* ≈ 320 × 10^−16^erg ≈ 1.9 kJ/mol, or *B* ≈ 0.85 *k*_*B*_*T*_0_ per one surface H_2_O molecule. Thus,

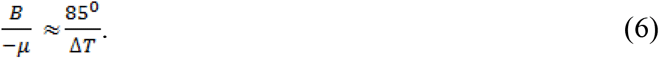

As a result, the transition state free energy for the pathway of a 3D ice crystal formation is estimated as

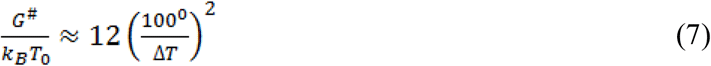

Then, according to the transition state theory(25, 26)^25,26^, the time of appearance of an ice nucleus around one given H_2_O molecule can be estimated as 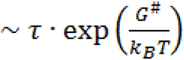, where τ is the time of addition of one more water molecule to the ice; it is no less than τ_0_∼10^−12^ s, the typical time of thermal vibrations at ≈273°K, or rather ∼10^−11^ s, the time of overturning (27) of an H_2_O molecule diffusing in the water at ≈273°К. More accurately, the value of *τ* is estimated as the difference between the rates of water attachment to and detachment from the ice (17): 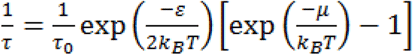, where *ε*≈51 kJ/mol is the energy of ice sublimation (23); at *T*≈273°К this gives

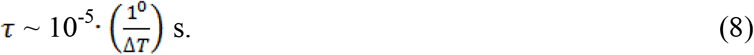

Since after the formation of the ice nucleus the remaining H_2_O molecules attach to it more or less independently, the *τ* value is both the time of attachment of one H_2_O molecule to the ice and the time of growth of one layer of H_2_O molecules upon the ice. The theoretical estimate (8) is in a good agreement with our experimental estimate of the time of growth of a layer of H_2_O molecules upon the ice: after the nucleation, 150 mg of H_2_O (i.e., ∼4.5·10^21^ molecules) form a piece of ice with ∼2·10^7^ H_2_O layers within ∼60 s at Δ*T* ≈ 5°.

If in the considered volume (e.g., in a cell) there are *N*_*V*_ water molecules, and a nucleus can arise around any of them, the characteristic time of appearance of one and only 3D ice nucleus in this volume can be estimated as

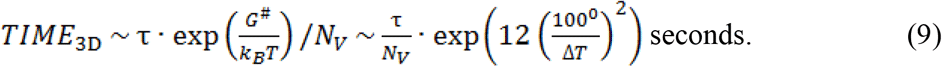

Now it is easy to find that

1. The smallest stable 3D pieces of ice surrounded by water (Fig. 3a) include *n*_*min*_ ≈ (5· 85° / Δ*T*)^3^ molecules (see Eqs. (3), (6)), that is, *n*_*min*_ ∼ 10^7^ H_2_O molecules at −2°C, *n*_min_ ∼ 24000 at −15°C, *n*_min_ ∼ 3000 at −30°C, and *n*_min_ ∼800 at −45°C, i.e., they are rather large, which justifies our neglect of the details of their structure.
2. Since *τ*∼10^−5^ -10^−6.7^ s at 1°-50°C (see Eq. (8)), in the volume of a typical animal cell (containing *N*_*V*_ ∼10^15^ H_2_O molecules), ice can appear (see Eq. (9) at −2°C in ∼10^13008^ seconds, i.e., ∼10^12999^ years (!!!), at −15°C – in ∼10^211^ seconds or ∼10^202^ years (!!!), at −30°C in ∼10^28^ years (!!!), and only at −45°C water in a cell can freeze in hours; that is, ice can practically never appear in pure bulk water even at −30°C (and not only in the volume of an animal cell but also in a swimming pool (28)), and this correlates with the observation that water droplets in the atmosphere freeze below -35°C only (29).

**Fig. 3.**
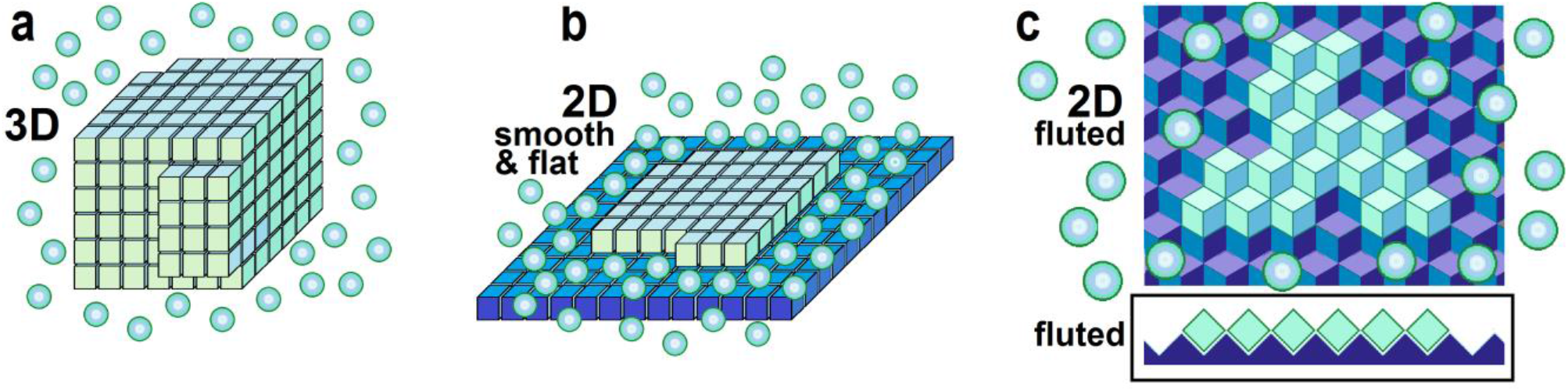
Schematic representations of three-dimensional (3D) and two-dimensional (2D) crystal nuclei. The 3D nucleus (**a**) appears in bulk liquid; 2D ice nuclei arise only on an ice or ice-binding surface that can be flat & smooth (**b**), fluted (**c**) or corrugated (see the inset to (**c**) that aims to illustrate that contacts between molecules within the “fluted” or “corrugated” layer are much smaller than contacts of molecules forming such a layer with molecules of the lower and higher layers (28)). The molecules of the crystal and layers are depicted in light-blue or greenish-blue cubes; molecules of the ice-binding surface in dark-blue or violet-blue cubes; “free” liquid molecules in balls. The additional free energy of a molecule of the surface of a 3D nucleus (**a**) or of the perimeter of a 2D nucleus on the smooth surface (**b**) is significant (see the text), while the additional free energy of a molecule of the perimeter of a 2D nucleus on the fluted (**c**) or corrugated surface is close to zero due to the small contact between molecules within such layers; therefore, they are less compact, but can form faster (see the text).

It’s another matter when ice arises not in bulk water but on some, even very small, ice-binding surface.

If such a surface can bind ice (according to Faraday, this property is possessed by flannel but not by gold (18)), then an ice nucleus arises on it not as a 3D but as a 2D object (Fig. 3b, 3c). We are not interested in the first monomolecular layer of ice on the intracellular surface: when the binding is strong, this monomolecular layer can also appear (but not grow further) at positive temperatures. We are interested in the beginning of further “growth of the ice layer on ice”, which can only occur at negative temperatures, leading to freezing of the cell. And although the formation of a two-dimensional layer is not, strictly speaking, the first-order phase transition, this layer is still rather similar to a crystal (30), and we can approximately estimate its nucleation using equations similar to those given above. When an ice layer freezes onto the previous layer of ice, the additional free energy associated with the boundary of the new layer depends only on its perimeter but not on the new layer’s area (as one ice surface in this case only replaces the other, see Figs. 3b, 3c).

The free energy of a 2D layer on the surface can be expressed in the same form that was already used in the 3D case,

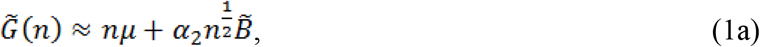

(with the accordingly changed power of *n*, of course), but the values *α*_2_ and 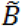 that describe the boundary free energy are now different from the values *α*_3_ and *B* used in the 3D case. Now, 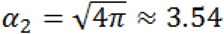 for a circular perimeter and *α*_2_ = 4 for a square perimeter, so that *α*_2_ ≈ 4 can be taken for our rough estimates. The value 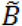 for a molecule of the perimeter of a smooth layer (Fig. 3b) should be close to the value of *B* for a molecule of the surface of a 3D body (Fig. 3a), but for a molecule of the perimeter of a fluted layer (Fig. 3c), the value of 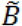 should be close to zero due to the small contact of molecules within the fluted (see Inset in Fig. 3c) layer.

The value of 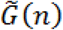 has its maximum 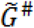 at *n* = *ñ*_0_, where 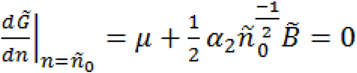, so that

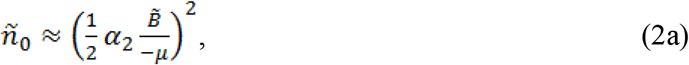

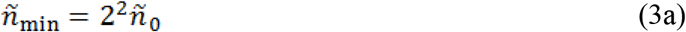

being the number of molecules in the minimally stable 2D piece (“seed”) of ice, and

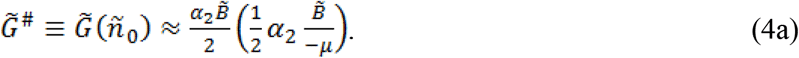

With 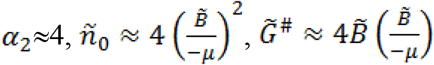, the minimally stable 2D layer of ice contains

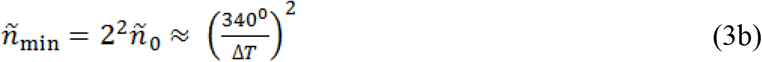

particles if the layer is smooth and thus 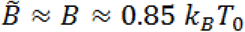; if the layer is fluted or corrugated (as in Fig. 3c), and thus 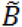 is close to zero, then *ñ*_0_, *ñ*_min_ and 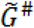 are close to zero too.

In other words, on fluted ice-binding surfaces ice can arise non-cooperatively, without nucleation of a phase transition, and therefore without delay. Non-cooperative formation of ice may also occur on an ice-binding surface, which is bent like the inner surface of a gutter or cube (for details, see Ref. 28). In these cases, one can expect fast freezing just at 0°C, without overcooling of the liquid.

At the same time, ice formation on flat and smooth (shown in Fig. 3b) surfaces, like the ice formation in bulk water, occurs at sub-zero temperatures, with a large (and sometimes enormous, see below) delay caused by the need to form an ice crystal nucleus; it is this process of nucleation of a smooth, not fluted layer that remains for us to consider.

As a result of nucleation at the smooth ice-binding surface (when 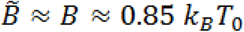), the free energy of the transition state on a pathway to the ice crystal formation is estimated as

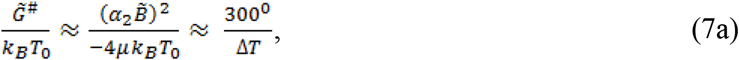

and the characteristic time of appearance of a 2D ice nucleus on the smooth surface capable of accommodating *N*_*S*_ molecules of H_2_O is estimated (given that 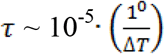 seconds, see Eq. (8)) as

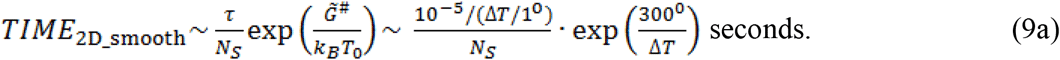

If all *N*_*V*_ ∼10^15^ water molecules in the cell are adjacent to a smooth (with 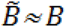) ice-binding surface (i.e., *N*_*S*_ = *N*_*V*_), then a 2D ice nucleus appears on such a surface at Δ*T*=2° (i.e., at −2°C), within ∼10^45^ seconds (≈10^36^ years (!!!)), and at −4°C – within ∼10^12^ seconds or ≈3000 years, i.e., ice will never appear within a living cell at −4°C, though it has a chance to appear at −4°C within a blood vessel whose surface is many millions of times larger than that of a cell.

However, already at −5°C, an ice nucleus can appear not only within a blood vessel, but also on a cell-size ice-binding surface in a couple of days, and at −6°C – in just a few seconds.

If (which is more natural) only *N*_*S*_∼*N*_*V*_ ^2/3^ ∼10^10^ waters (out of *N*_*V*_∼10^15^ waters in the cell) are adjacent to the ice-binding smooth surface, then the 2D ice nucleus arises at such a surface at −5°C within ∼100 years (an age! – but this is for a cell; for a million times greater blood vessel this will take only an hour…) A 2D ice nucleus arises within ∼10 days at −6°C, but only some seconds at −8°C (it seems that our experiments, whose results are shown in Fig. 1 and Table 1, dealt with this type of ice formation). Note that the presence of dust or other microparticles in water may entail an increase of *N*_*S*_ by a factor of ∼100 and, accordingly, result in a 100-fold reduction of the nucleation time; but even without the dust, the nucleation will take milliseconds at −10°C, and a fraction of the microsecond at −15°C.

A tremendous acceleration of the appearance of an ice nucleus with even a slight decrease in temperature!

It should be noted that a fairly large (contacting with 10^10^ − 10^15^ waters, see above) ice-binding smooth surface is needed for the emergence of an ice nucleus in a reasonable time. The minimal total size of this surface — which can consist of several parts — can be estimated for given *TIME*_2D_smooth_ and Δ*T* values from Eq. (9a). However, the size of the minimally stable two-dimensional ice piece, *ñ*_min_, should not be too large (see Eq. (3b)): *ñ*_min_ ≈170^2^ waters at −2°C (diameter of such a piece of ice is ≈60 nm), *ñ*_min_ ≈70^2^ at −5°C (diameter – ≈25 nm), *ñ*_min_ ≈40^2^ at −8°C (diameter – ≈15 nm), and *ñ*_min_≈22^2^ at −15°C (diameter – ≈8 nm).

So far, we took into account only the time of initiation of ice formation, but not the time of its growth. The reason is that growth is a relatively fast process. The time of addition of one new layer of water molecules to the ice at temperatures below −1°C is less than 10^−5^s (see Eq. (8)). So, during 1 second, 100000 new layers of H_2_0 molecules join the ice (this new layer is ∼30 μm thick like a living cell), – while, speaking on the ice initiation, we spoke about much longer times. Thus, the time of ice growth can be neglected.

The above estimates show that ice-binding surfaces of sufficient size and kinetic phenomena associated with the appearance of the ice nuclei are drastically important for the freezing.

The ice-binding properties of various surfaces, mainly of technical use, have been studied (see, e.g., Refs. (18), (31), (32) and the literature in Refs. (31, 32). However, we know practically nothing about the ice-binding properties of significant surfaces of biological origin (which can be the targets for IBPs); thus, the identification of such surfaces and the study of their properties will be the next step in the investigation of the action of antifreeze proteins.

Returning to experiments on the freezing solutions, and seeing (Fig. 1 and Table 1) that all of them freeze within minutes at -7.9°C ÷ -14.7°C, we can conclude that: (1) The ice nucleus appears not in bulk solution (where even thousands of years would not be enough for this), and not on some fluted or bent surfaces (otherwise, freezing would occur quickly, without preliminary overcooling of the liquids), but somewhere on relatively smooth walls of the test tubes or dust particles; (2) Judging from the degree and time of the overcooling, the ice-binding surfaces within our test tubes (estimated from Eq. (9a)) covered less than 10 μm^2^; (3) Since the addition of IBP (that can bind to ice, thus decreasing the 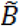 value) raises the freezing temperature by no more than ∼1°C (i.e., by no more than ∼10%, see Fig. 1), the IBP-induced decrease of the 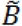 value (which can be estimated using Eqs. (7a), (9a)) is minor, about 5% or less.

Returning to the ice-binding proteins: if the cell or blood vessel has significant (∼μm^2^ in total) ice-binding surfaces, some kind of antifreeze molecules are really needed to bind to these surfaces and block the growth of ice there.

Now, let us see if cells do have surfaces that bind the ice-binding proteins.

To verify this hypothesis, we constructed a gene of the fusion protein GFP-mIBP83 from genes of the green fluorescent protein (33) (GFP) and the ice-binding protein mIBP83, (see Ref.(15)) which is a mutant of cfAFP antifreeze protein (12, 13). The mutant form of GFP, called “cycle3 GFP” was chosen for the experiments because we have studied it previously and found that it remains monomeric and fluorescent under native conditions (34) (hereafter, it is called “GFP”). The fusion protein was expressed in *E*.*Coli* cells, isolated, and purified (15).

A simple experiment has demonstrated that mIBP83 connected with GFP retains the ability of binding to the ice surface.

The left part of Figure 4 shows a test tube marked “+IBP” with ice floating in a solution of GFP-mIBP83 illuminated by UV light that excites the fluorescence of GFP. Here, one can see the contours of luminous ice because the protein GFP-mIBP83 has covered the ice surface. A control experiment was performed using a GFP solution. The right part of Figure 4 shows a test tube marked “-IBP” with ice floating in a solution of the GFP protein. Here, one can see that mainly the solution glows, while the contours of the ice are not visible (for more photos, see) (15).

**Fig. 4.**
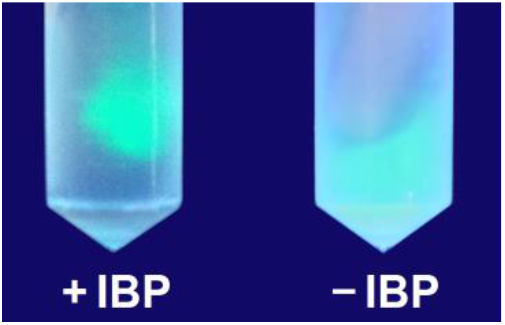
Experiments on the IBP-to ice-binding. Comparison of the test-tubes with pieces of ice in solutions of GFP-mIBP83 (+IBP) and solely GFP (-IBP) shows that GFP-mIBP83 binds to the ice, while GFP does not.

The obtained fused protein GFP-mIBP83 allowed visualization of intracellular localization of IBP. For this purpose, the human breast cancer cells SKBR-3 were transfected using plasmid vectors encoding either the fused protein GFP-mIBP83 or solely GFP. The transfected cells were cultured under standard conditions. To test the response to cold, the cells were incubated first at +37°C, then at +2°C for 2 hours and immediately fixed with 4% formaldehyde to prevent protein redistribution during the imaging procedure. At lower temperatures, the cells separated from the substrate and were inconvenient for research (for details, see *Cell culture experiments*).

The pattern of intracellular localization of GFP-mIBP83 does differ from that of the sole GFP just at low positive (+2°C) temperatures (Figure 5). At +37°C, both proteins are mainly distributed between the cytoplasm and the nucleus. At +2°C, the stained GFP-mIBP83 is concentrated in small vesicles or granules which can be clearly seen in the cells. This suggests that GFP-mIBP83 specifically binds to some cellular structures at cooling to zero temperature. One can see that this cooling entails drastic changes in GFP-mIBP83 distribution, as compared to that at +37°C. The amount of diffusely distributed protein decreased, and GFP-mIBP83 accumulated in perinuclear regions. These regions are known to be rich in cellular organelles, such as cisternae of the Golgi apparatus, mitochondria, endoplasmic reticulum, late endosomes, etc. Of course, our next goal will be a detailed investigation of the targets of IBP binding.

**Fig. 5.**
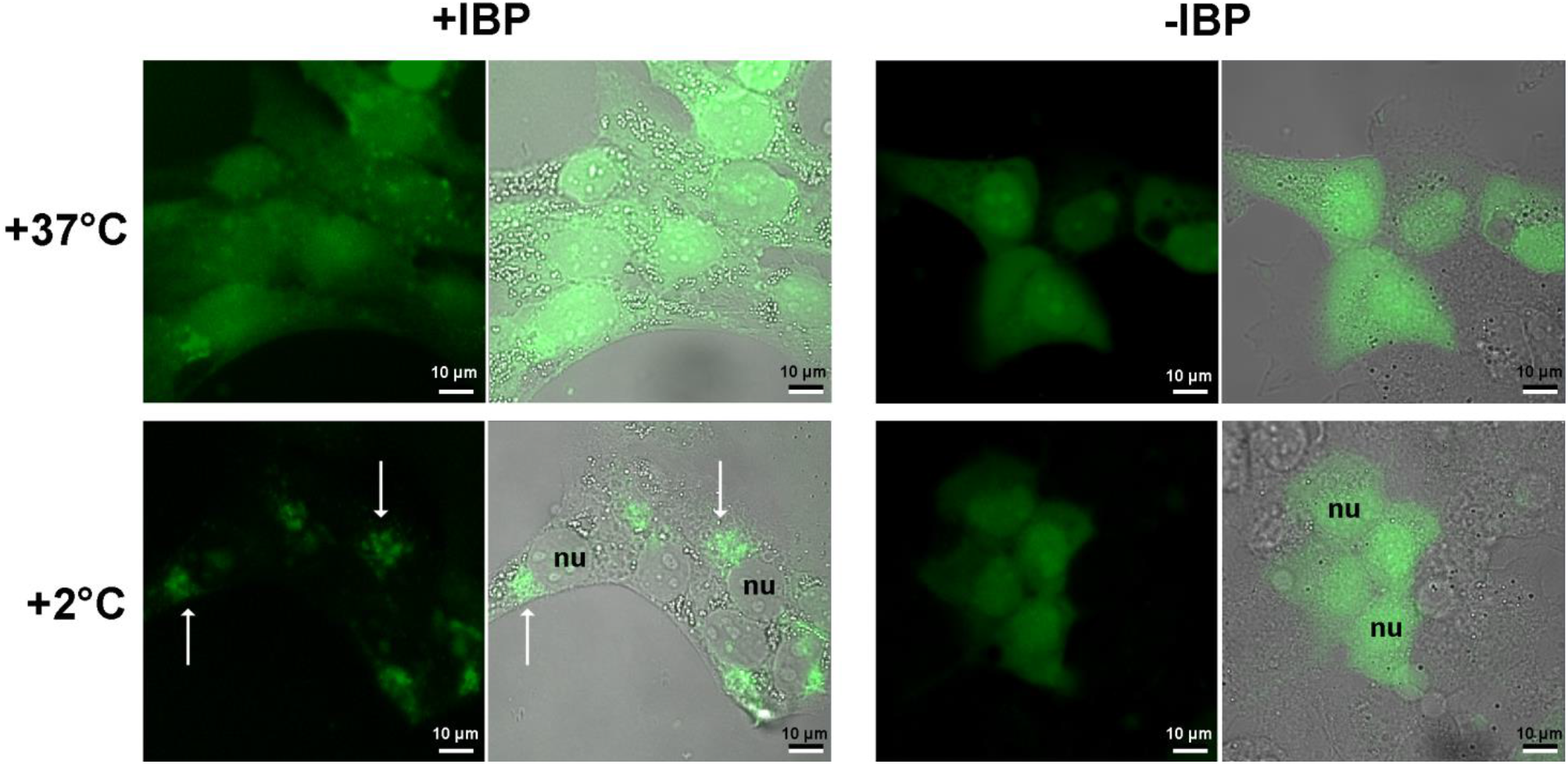
Localization of the fused protein GFP-mIBP83 and the solely GFP in SKBR-3 cells. The cells were kept at +37°C or incubated at +2°C for 2 h, then fixed and imaged using a laser scanning microscope. The fluorescence images and the merged transmittance and fluorescence images are presented for each variant of treatment. The nuclei (**nu**) of individual cells are marked. The white arrows indicate the most pronounced GFP-mIBP83 accumulations in the perinuclear regions of cooled cells. Bar, 10 µm. (For details, see *Cell culture experiments*).

The obtained results are in line with our hypothesis that, at temperatures of about 0°C, the cell contains some ice-nucleating surfaces to which GFP-mIBP83 can bind while GFP cannot. On such surfaces, ice nuclei can arise at low temperatures if these surfaces are not blocked by IBPs.

It should be stressed that in our experiments, at +2°C, GFP-mIBP83 binds to the cell elements directly, and not via ice, that cannot form at +2°C.

Since mIBP83 binds the ice *in vitro* (Fig. 4), it can stabilize the ice and so it does indeed (see Fig. 2); thus, an IBP can serve not only as antifreeze but also as an ice-nucleating protein. However, as follows from the above calculations, the linear size of an ice-nucleating surface must be not less than ∼10-20 nm. This accords to the statement of Bissoyi at al. that large (∼200 kDa) IBPs can initiate the formation of ice nuclei (35) while smaller IBPs serve only as antifreeze that prevents the growth of ice crystals.

The IBP binding to cellular surfaces should stabilize these surfaces, and thus it can protect cells from the hypothermic cold shock (preceding the emergence of ice) damage; this is consistent with the hypothesis by Hirano et al. (6).

Thus, the proposed binding of IBPs to cellular surfaces can simultaneously explain both phenomena experimentally observed (10, 35) upon cell freezing: (i) the tolerance of cells to the cold shock under moderate cooling to 0° (possibly caused by the stabilization of cell surfaces caused by their binding to IBPs), and (ii) survival of cells upon stronger freezing (caused by the IBP-induced inhibition of the ice crystal appearance).

It is worth emphasizing that this work gives a fundamentally different view on the functioning of “ice-binding” (or “antifreeze”) proteins. Their task is not to bind to ice crystals and stop their growth but to bind – directly or through the thinnest layer of water molecules – to those cell or tissue surfaces where ice nuclei can form, thus preventing the ice formation altogether. A blockage of these ice-nucleating surfaces is the only way to prevent freezing using a small number of antifreeze molecules.

## Freezing and melting experiments equipment

Experiments on melting and freezing were carried out using a thermostat Julabo F-25, Germany. The thermostat and measuring thermometers were checked using a laboratory thermometer LT-300-N, TERMEX, Russia, resolution 0.01°C, accuracy ±0.05°C.

The test tubes (shown in figure 4) were illuminated using a transilluminator ecx-f20.m VILBER, France.

## Cell culture experiments

The human breast adenocarcinoma cells SKBR-3 (ATCC^®^ HTB-30™) were cultured in the McCoy’s medium (PanEco, Russia) with 10% (v/v) fetal calf serum (HyClone, USA) in 5% CO_2_ at 37°C.

For transient expression of fluorescent proteins, we used the plasmid vectors pTag-2N encoding the gene of GFP-mIBP83 or sole GFP (cycle3 GFP) under the control of cytomegalovirus promoter and the gene of resistance to the antibiotic G418. The cells were transfected using Lipofectamin 3000 (Invitrogen, USA) according to the manufacturer’s instructions, followed by cultivation in selective G418-containing medium for several passages.

Cooling of the cells was performed using a solid-state ThermoStat Plus (Eppendorf^®^, Germany) with precise temperature control. The cells were cultured in Falcon^®^ 96-well black/blear flat-bottom TC-treated imaging microplates or Eppendorf^®^ glass-bottom cell imaging dishes. To test the response to cold, the cell cultures were incubated at +2°C for 2 h and immediately fixed with 4% formaldehyde. The temperature of +2°C was chosen as the lowest temperature at which the cells remained spread out, attached to the substrate, and accordingly, convenient for research.

The images were obtained using a laser scanning microscopy system Axio Observer Z1 LSM-710 DUO NLO (Carl Zeiss, Germany). The GFP fluorescence was excited at 488 nm and registered in a wide spectral range of 500-735 nm.

## Acknowledgements

We are grateful to G. Fermi, A. Yu. Grosberg, M. D. Frank-Kamenetsky, E. G. Malenkov, E. I. Shahnovich, D. Fraenkel, and A. N. Gavrishev for fruitful discussions, and to G. S. Nagibina, Yu. D. Okulova, E. N. Samatova, and E. V. Serebrova for assistance. The work was supported by the RAS Program №9 on Fundamental Research, grant № 01201358029, and by the Russian Foundation for Basic Research grant No. 19–04–00420.

## Author Contributions

A.V.F. and B.S.M. conceived the idea for the study, participated in experimental design, performed data analysis, and wrote the paper; K.A.G - plasmids construction; B.S.M., E.A.S., and I.V.B. performed the experiments.

## Author Information

The authors declare no competing financial interests. Readers are welcome to comment on the online version of the paper. Correspondence and requests for materials should be addressed to B.S.M.

## References

1. P. A. Cziko, A. L. DeVries, C. W. Evans, C.-H. C. Cheng, Antifreeze protein-induced superheating of ice inside Antarctic notothenioid fishes inhibits melting during summer warming. Proc. Natl. Acad. Sci. 111, 14583–14588 (2014).

2. P. F. Scholander, L. van Dam, J. W. Kanwisher, H. T. Hammel, M. S. Gordon, Supercooling and osmoregulation in arctic fish. J. Cell. Comp. Physiol. 49, 5–24 (1957).

3. A. L. DeVries, Glycoproteins as Biological Antifreeze Agents in Antarctic Fishes. Science (80-.). 172, 1152–1155 (1971).

4. A. L. DeVries, Biological antifreeze agents in coldwater fishes. Comp. Biochem. Physiol. Part A Physiol. 73, 627–640 (1982).

5. R. Surís-Valls, I. K. Voets, The Impact of Salts on the Ice Recrystallization Inhibition Activity of Antifreeze (Glyco)Proteins. Biomolecules 9, 347 (2019).

6. Y. Hirano, et al., Construction of Time-Lapse Scanning Electrochemical Microscopy with Temperature Control and Its Application To Evaluate the Preservation Effects of Antifreeze Proteins on Living Cells. Anal. Chem. 80, 9349–9354 (2008).

7. A. Jorov, B. S. Zhorov, D. S. C. Yang, Theoretical study of interaction of winter flounder antifreeze protein with ice. Protein Sci. 13, 1524–1537 (2004).

8. A. L. DeVries, Survival at freezing temperatures. Biochem. Biophys. Perspect. Mar. Biol., 289–330 (1974).

9. Y. Celik, et al., Microfluidic experiments reveal that antifreeze proteins bound to ice crystals suffice to prevent their growth. Proc. Natl. Acad. Sci. 110, 1309–1314 (2013).

10. M. Kuramochi, et al., Expression of Ice-Binding Proteins in Caenorhabditis elegans Improves the Survival Rate upon Cold Shock and during Freezing. Sci. Rep. 9, 6246 (2019).

11. D. Doucet, et al., Structure-function relationships in spruce budworm antifreeze protein revealed by isoform diversity. Eur. J. Biochem. 267, 6082–6088 (2000).

12. E. K. Leinala, et al., A β-Helical Antifreeze Protein Isoform with Increased Activity. J. Biol. Chem. 277, 33349–33352 (2002).

13. E. K. Leinala, P. L. Davies, Z. Jia, Crystal Structure of β-Helical Antifreeze Protein Points to a General Ice Binding Model. Structure 10, 619–627 (2002).

14. M. G. Tyshenko, D. Doucet, P. L. Davies, V. K. Walker, The antifreeze potential of the spruce budworm thermal hysteresis protein. Nat. Biotechnol. 15, 887–890 (1997).

15. K. A. Glukhova, J. D. Okulova, B. S. Melnik, Designing and studying a mutant form of the ice-binding protein from Choristoneura fumiferana. BioRxiv (2020).

16. A. R. Ubbelohde, Melting and crystal structure. Chapter 14. (Clarendon Press, Oxford, 1965).

17. A. A. Chernov, Modern Crystallography III vol. 36 (Springer Berlin Heidelberg, 1984).

18. M. Farraday, Experimental Researches in Chemistry and Physics. Chapters “On ice and freezing water”, pp. 372–377, and “On Regelation”. pp. 377–382. (Taylor and Francis, London, 1859).

19. J. J. Métois, J. C. Heyraud, The overheating of lead crystals. J. Phys. 50, 3175–3179 (1989).

20. M. Forsblom, G. Grimvall, How superheated crystals melt. Nat. Mater. 4, 388–390 (2005).

21. Q. S. Mei, K. Lu, Melting and superheating of crystalline solids: From bulk to nanocrystals. Prog. Mater. Sci. 52, 1175–1262 (2007).

22. W. J. Smit, H. J. Bakker, The Surface of Ice Is Like Supercooled Liquid Water. Angew. Chemie Int. Ed. 56, 15540–15544 (2017).

23. D. R. Lide, CRC Handbook of Chemistry and Physics on CD (Section 6) (CRC Press, Boca Raton, 2005).

24. W. B. Hillig, Measurement of interfacial free energy for ice/water system. J. Cryst. Growth 183, 463–468 (1998).

25. N. M. Knorre, D.G., Emanuel, The Course of Chemical Kinetics. 4th Ed. Chapters II, III, V. (Higher Sch. Moscow, 1984) (1984).

26. H. Eyring, The Activated Complex in Chemical Reactions. J. Chem. Phys. 3, 107–115 (1935).

27. A. V. Finkelstein, O. B. Ptitsyn, Protein Physics. A Course of Lectures, 2nd edition. Lecture 8. (Academic Press, An Imprint of Elsevier Science, Amsterdam • Boston • Heidelberg • London • New York • Oxford • Paris • San Diego •San Francisco • Singapore • Sydney • Tokyo; 2016).

28. A. V Finkelstein, Some peculiarities of water freezing at small sub-zero temperatures. Preprint. ArXiv (2020).

29. T. Koop, B. J. Murray, A physically constrained classical description of the homogeneous nucleation of ice in water. J. Chem. Phys. 145, 211915 (2016).

30. K. J. Strandburg, Two-dimensional melting. Rev. Mod. Phys. 60, 161–207 (1988).

31. Raraty, The adhesion and strength properties of ice.

32. Work, A critical review of the measurement of ice adhesion to solid substrates.

33. H. Fukuda, M. Arai, K. Kuwajima, Folding of green fluorescent protein and the cycle3 mutant. Biochemistry 39, 12025–12032 (2000).

34. K. F. Glukhova, V. V Marchenkov, T. N. Melnik, B. S. Melnik, Isoforms of green fluorescent protein differ from each other in solvent molecules “trapped” inside this protein. J. Biomol. Struct. Dyn. 35, 1215–1225 (2017).

35. A. Bissoyi, et al., Ice Nucleation Properties of Ice-binding Proteins from Snow Fleas. Biomolecules 9, 532 (2019).

